# Thg1 family 3’-5’ RNA polymerases as tools for targeted RNA synthesis

**DOI:** 10.1101/2024.02.24.581873

**Authors:** Malithi I. Jayasinghe, Krishna J. Patel, Jane E. Jackman

## Abstract

Members of the 3’-5’ RNA polymerase family, comprised of tRNA^His^ guanylyltransferase (Thg1) and Thg1-like proteins (TLPs), catalyze templated synthesis of RNA in the reverse direction to all other known 5’-3’ RNA and DNA polymerases. Discovery of enzymes capable of this reaction raised the possibility of exploiting 3’-5’ polymerases for post-transcriptional incorporation of nucleotides to the 5’-end of nucleic acids without ligation, and instead by templated polymerase addition. To date, studies of these enzymes have focused on nucleotide addition to highly structured RNAs, such as tRNA and other non-coding RNA. Consequently, general principles of RNA substrate recognition and nucleotide preferences that might enable broader application of 3’-5’ polymerases have not been elucidated. Here, we investigated the feasibility of using Thg1 or TLPs for multiple nucleotide incorporation to the 5’-end of a short duplex RNA substrate, using a templating RNA oligonucleotide provided *in trans* to guide 5’-end addition of specific sequences. Using optimized assay conditions, we demonstrated a remarkable capacity of certain TLPs to accommodate short RNA substrate-template duplexes of varying lengths with significantly high affinity, resulting in the ability to incorporate a desired nucleotide sequence of up to 8 bases to 5’-ends of the model RNA substrates in a template-dependent manner. This work has further advanced our goals to develop this atypical enzyme family as a versatile nucleic acid 5’-end labeling tool.

## INTRODUCTION

Canonical DNA and RNA polymerases incorporate nucleotides in the 5’-3’ direction by virtue of their conserved reaction chemistry. For these enzymes, the terminal 3’-hydroxyl of a polynucleotide chain performs a nucleophilic attack on the α-phosphate of the incoming 5’-triphosphorylated nucleotide (NTP), releasing pyrophosphate and resulting in new phosphodiester bond formation at the 3’-end of the growing nucleic acid (Joyce and Steitz 1995; Steitz 1998; Steitz 1999). Nucleotide polymerization can occur in the opposite 3’-5’ direction using the same chemistry when the orientation of the substrates is reversed. For this reaction, the 3’-hydroxyl of the incoming NTP performs the nucleophilic attack on the α-phosphate at the 5’-end of a growing 5’-triphosphorylated polynucleotide, also resulting in the formation of a standard phosphodiester bond, but with the new NTP added at the 5’-end. The enzyme family comprised of tRNA^His^ guanylyltransferase (Thg1) and Thg1-like proteins (TLPs) provided the first examples of enzymes capable of performing 3’-5’ polymerase chemistry, and roles for these enzymes in tRNA processing and repair has now been established for family members in multiple species (Jackman and Phizicky 2006b; Abad et al. 2010; Abad et al. 2011; Jackman et al. 2012; Long et al. 2016; Chen et al. 2019).

The founding enzyme of the Thg1/TLP family, *Saccharomyces cerevisiae* Thg1 (ScThg1), was discovered because of its ability to incorporate a single essential G nucleotide (G_-1_) at the 5’-end of tRNA^His^ (Cooley et al. 1982; Gu et al. 2003; Gu et al. 2005). Thg1-type enzymes are broadly conserved in Eukarya where their essential role in tRNA^His^ processing is likely highly conserved, and consequently these enzymes exhibit strong selectivity for tRNA^His^ over other tRNA species (Rao and Jackman 2015; Edvardson et al. 2016; Long et al. 2016). However, TLP-type enzymes are found in all three domains of life and appear to play more diverse physiological roles that predictably necessitate recognition of a broader set of RNA substrates for 3’-5’ addition (Jackman et al. 2012; Chen et al. 2019). The eukaryotic slime mold *Dictyostelium discoideum* encodes three different TLPs (DdiTLP2, DdiTLP3 and DdiTLP4) that exhibit non-redundant functions *in vivo* (Abad et al. 2011; Long et al. 2016). While DdiTLP2 incorporates G_-1_ selectively into mitochondrial tRNA^His^, DdiTLP3 repairs 14 different 5’-truncated tRNAs during a process known as mitochondrial tRNA 5’-editing (**Figure 1**)(Long and Jackman 2015; Long et al. 2016). Interestingly, although DdiTLP4 is essential for growth in *D. discoideum*, its biological function remains unknown. Other protozoan TLPs likely use 3’-5’ polymerase activity to repair 5’-truncated tRNAs similar to DdiTLP3, since mitochondrial 5’-editing of tRNA appears to be widespread in protozoa. The amoebae *Acanthamoeba castellanii* was the first organism in which tRNA 5’-editing was described and indeed encodes two TLPs (AcaTLP1 and AcaTLP2) (Lonergan and Gray 1993; Laforest et al. 1997; Laforest et al. 2004; Betat et al. 2014). Although a role in tRNA 5’-editing has not been demonstrated conclusively for either *A. castellanii* enzyme, both enzymes catalyze 5’-end repair of predicted tRNA editing intermediates *in vitro* (Rao et al. 2013). Notably, TLP enzymes are also found in bacterial and archaeal species where a process similar to mitochondrial 5’-editing of tRNA does not appear to occur, and therefore other functions for these enzymes are likely, and remain to be demonstrated (Abad et al. 2010; Rao et al. 2011; Heinemann et al. 2012; Kimura et al. 2016; Chen et al. 2019).

**Figure 1:**
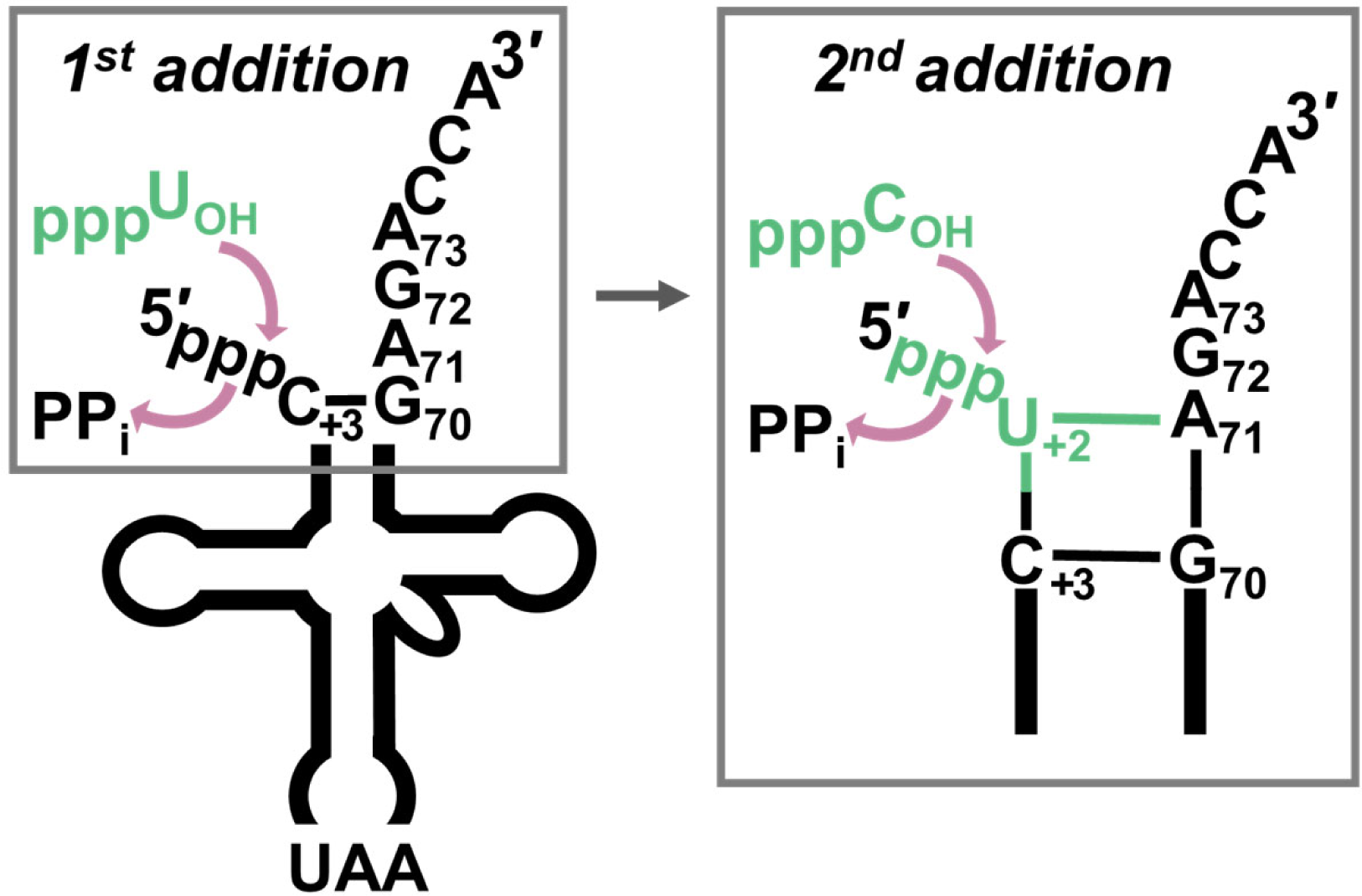
3’-5’ polymerase activity is used to repair mitochondrial tRNA. A schematic of tRNA 5’-end repair during mitochondrial tRNA editing in protozoa. The *D. discoideum* mitochondrial tRNA^Leu^ substrate shown in the diagram corresponds to an intermediate generated during 5’-editing that is missing nucleotides U_+2_ and C_+1_ from its 5’-end. In *D. discoideum*, DdiTLP3 utilizes the A_71_ and G_72_ nucleotides of the tRNA 3’-acceptor stem as a template for selection and incorporation of U_+2_ and C_+1_, respectively, via Watson-Crick dependent 3’-5’ polymerase activity. In the box showing the first (U_+2_) nucleotide addition, the 3’-OH of an incoming UTP (green) performs a nucleophilic attack on the α-phosphate of the 5’-triphosphorylated end of the truncated RNA. Likewise, during the second (C_+1_) addition, an incoming CTP uses its 3’-OH to attack the α-phosphate of the added U_+2_ nucleotide, thus restoring the fully base-paired aminoacyl-acceptor stem on the tRNA that is required for translation.

Consistent with the distinct biological roles for Thg1 vs. TLP enzymes, different members of the 3’-5’ polymerase family exhibit distinct *in vitro* biochemical properties. Thg1 is strongly selective for activity on its physiological substrate tRNA^His^ over other tRNAs, which is accomplished by recognition of the unique GUG anticodon sequence in tRNA^His^ (Jackman and Phizicky 2006a). Alternatively, nearly all TLPs tested to date catalyze tRNA repair (as in Figure 1) on a broad variety of tRNA substrates *in vitro*, and moreover exhibit a strong preference for incorporating multiple Watson-Crick base-paired nucleotides during these reactions (Abad et al. 2011; Rao et al. 2011; Rao et al. 2013; Desai et al. 2018). Since TLPs overall exhibit more flexible RNA substrate recognition than Thg1-type enzymes, we hypothesized that TLPs would be attractive candidates for harnessing 3’-5’ polymerase activity to develop a versatile nucleic acid 5’-end labeling tool. However, these efforts would be facilitated by a deeper understanding of RNA substrate and nucleotide preferences of TLPs outside of the context of the highly structured tRNA substrates that have so far been used in enzymatic characterization of 3’-5’ polymerases(Rao et al. 2011; Smith and Jackman 2012; Desai et al. 2018; Patel et al. 2021). Here we describe the first analysis of template-dependent 3’-5’ polymerase activities catalyzed by TLPs with model RNA substrates using an *in vitro* gel-based 5’-extension assay. Two different TLP enzymes (AcaTLP2 and DdiTLP4) robustly incorporate 5’-nucleotides into a short (26-nucleotide) RNA substrate in the presence of an RNA oligonucleotide that serves as a template for addition. We demonstrate that these two TLPs efficiently incorporate specific nucleotides into the tested model RNA substrate in a template-dependent manner and can accommodate RNA duplexes with base-paired lengths from 7-19 base pairs with only moderate differences in catalytic efficiency. Biochemical characterization revealed that the ability of the most active TLPs to act on the model duplexes was driven by their unusually high binding affinity for the RNAs, in contrast to the binding affinities that are normally exhibited for tRNA by members of the Thg1/TLP family for highly structured tRNAs.

## MATERIALS AND METHODS

### Thg1/TLP protein expression and purification

Plasmids for expression and purification of ScThg1, MsTLP, BtTLP, MxTLP, AcaTLP2, DdiTLP3 and DdiTLP4 enzymes with N-terminal His6-tag sequences have been described previously(Jackman and Phizicky 2006b; Abad et al. 2010; Abad et al. 2011; Rao et al. 2011). Briefly, enzymes were expressed in *E. coli* BL21 (DE3) pLysS or Rosetta pLysS cells and overexpressed proteins were purified using immobilized metal-ion affinity chromatography (IMAC) with TALON resin (Clontech) (Abad et al. 2010). Proteins were dialyzed after elution from the column in a dialysis buffer containing 20 mM Tris pH 7.5, 500 mM NaCl, 4 mM MgCl_2_, 50% Glycerol, 1 μM ethylenediaminetetraacetic acid (EDTA) pH 8.0 and 1 mM DTT and stored at -20°C. Purity of the proteins were tested by SDS-PAGE and assessed to be >95 % pure. The BioRad protein assay was used to determine the concentrations of purified proteins for enzymatic assays described below.

### *In vitro* transcription of RNA substrates and templates

*RNA substrate (26-mer):* Two different *in vitro* transcription strategies were used to create the model 26-mer RNA substrate (5’-GGGAGACCACAACGGUUUCCCUCUAG-3’) used for this work. Initially, a pET-derived vector that is normally used for T7 RNA polymerase-dependent protein expression (AVA421) was linearized by digestion with XbaI (NEB) in a reaction containing 1X CutSmart buffer and 0.5 U/μl XbaI to be used for run-off transcription using T7 RNA polymerase to transcribe the first 26 nucleotides following the T7 promoter sequence to generate the standard model RNA substrate. However, later efforts to improve resolution of the gel-based assay employed a construct with a self-cleaving HDV ribozyme sequence added immediately 3’-to the end of the 26-mer sequence to avoid 3’-end heterogeneity in the resulting transcripts. The HDV sequence was cloned into the XbaI/EcoRV sites of the AVA421 vector to enable linearization for run-off transcription by digestion with EcoRV-HF (NEB) in a reaction containing 1X CutSmart buffer and 0.4 U/μl EcoRV-HF.

In either case, uniformly labeled RNA substrates were generated by *in vitro* transcription in 80 μl reactions that included 15 μg linearized DNA template and 10-50 μg T7 RNA polymerase in the presence of 50 mM Tris pH 8.0, 1 mM Spermidine, 30 mM MgCl_2_, 7.5 mM DTT, 1 μg inorganic pyrophosphatase (Roche), 4 mM CTP/GTP/UTP, 2 mM ATP and α-^32^P-[ATP] (Perkin Elmer). The linearized DNA template was then removed by adding 100 U of DNase I (Roche) to the transcription reaction and incubating at 37 °C for 30 min. The resulting labeled transcripts were gel-purified from a 20% polyacrylamide 8 M urea gel by incubating the crushed gel slice containing the 26-mer band in RNA elution buffer (0.5 M ammonium acetate, 0.1% SDS, 5 mM EDTA) overnight at 37 °C. The eluted RNA was further purified by phenol: chloroform: isoamyl alcohol (PCA 25:24:1; v:v:v) extraction followed by ethanol precipitation and resuspension in ddH_2_O. All RNA transcripts were quantified by scintillation counting and stored at -20 °C. The same procedure was used to transcribe an unlabeled 26-mer RNA substrate for use in as a competitor, except that the RNA concentration was quantified by A_260_ measurement.

*RNA templates*: *In vitro* transcription of RNA templates was performed using DNA duplex templates containing a DNA oligonucleotide corresponding to the T7 promoter sequence and a second DNA oligonucleotide that creates the complementary region (**Supplementary Table 1**) for the T7 promoter sequence as well as a 5’-overhang sequence to direct production of the desired RNA templates of varied lengths. T7 promoter and templating oligonucleotides were mixed to a final concentration of 25 μM and annealed by heating to 95 °C for 5 min followed by rapid cooling on ice for 15 min. These 25 μM duplex stocks were used at a final concentration of 2.5 μM in *in vitro* transcription reactions containing 50 mM Tris pH 8.0, 1 mM Spermidine, 20 mM MgCl_2_, 10 mM DTT, 4 mM each NTP, 2.5 μg inorganic pyrophosphatase (Roche) and ∼50 μg T7 RNA polymerase. A DNase I (Roche) treatment was performed, and the transcripts were gel-purified as described for the RNA substrate transcription reactions above.

Template transcripts were treated with calf intestinal phosphatase (CIP) to remove 5’-phosphates that might enable them to be acted on by Thg1/TLPs, although even if this activity did occur on unlabeled template RNA, it would not be visualized in the assays. CIP reactions contained a final concentration of 1X NEB rCutSmart and 1 μl NEB quick CIP per 1 μg RNA and incubated at 37 °C for 30 min, followed by PCA extraction and ethanol precipitation before resuspending in 10-20 μl ddH_2_O.

### Template-dependent 5’-extension assay

RNA duplexes were created by annealing labeled 5’-triphosphorylated model RNA substrate (∼20 nM) to each RNA template (5 μM) by heating to 95 °C for 5 min followed by slow cooling to room temperature. The annealed duplexes were added to assay buffer containing 25 mM HEPES pH 7.5, 10 mM MgCl_2_, 3 mM DTT, 25 mM NaCl and 0.2 mg/ml BSA and the indicated NTPs (0.5 mM each). Reactions were initiated by addition of Thg1/TLP enzyme (final concentration 15 μM) and incubated at 27 °C for 2 hours. The reactions were quenched by addition of 100 mM EDTA to a final concentration of 10 mM. To each quenched reaction, unlabeled substrate RNA was added to act as a competitor for binding to the RNA template (5 μM final concentration, unless otherwise indicated in the assay), followed by addition of DMSO to a final concentration of 42.8% DMSO. Finally, an equal volume of formamide loading dye was added (10 ml of 100% formamide, 100 μl of 0.5 M EDTA, 1 μl of 10% bromophenol blue and 1 μl of 10% xylene cyanol), followed by heat denaturation at 95 °C for 5 min. Reactions were resolved using 20% polyacrylamide (19:1)/8M urea gels. The gels were visualized by Typhoon Trio Imager and quantified using ImageQuant software.

The same assay conditions were used to calculate single turnover rates of multiple nucleotide incorporation (*k_obs_*) at saturating concentrations of NTPs (0.5 mM) and saturating enzyme concentration (15 μM). At each time point, a 2 μl aliquot was taken out and mixed with EDTA to quench the reactions prior to gel resolution as described for end point assays above. Equation 1 was used to calculate *k_obs_* at each enzyme concentration using Kaleidagraph (Synergy software), where % product at each time point (P_t_) was plotted vs. time and fit to the single exponential rate equation, where ΔP is maximal product formation:

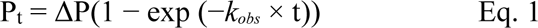

### Filter binding assay

RNA duplexes were created by mixing the uniformly labeled 5’-triphosphorylated model RNA substrate with equimolar amounts of the corresponding RNA template and heating to 95 °C for 5 min followed by slow cooling to room temperature to anneal. Binding reactions were performed in 50 μl reactions containing 1 nM RNA duplex, 25 mM HEPES pH 7.5, 10 mM MgCl_2_, 3 mM DTT and 125 mM NaCl. After addition of 0.1-10 μM purified Thg1/TLP enzyme, reactions were incubated at 27 °C for 45 min. For each reaction, 45 μl was spotted to a double-membrane system on a dot-blot apparatus with an upper nitrocellulose membrane and a lower hybond-N^+^ membrane (GE Healthcare). After filtration under vacuum, membranes were dried and imaged using a Typhoon imager, and protein-bound and free RNA were quantified from nitrocellulose and hybond membranes, respectively, using ImageQuant. The average fraction bound (F) for each reaction was calculated from reactions performed in triplicate and plotted against the enzyme concentration ([E]), with Eq. 2 used to calculate *K_D_* for each enzyme-RNA pair using Kaleidagraph (Synergy software). Eq. 2 was used to calculate *K_D_* from the plot of % bound vs. enzyme concentration [E] where *K_D_* is the dissociation constant, and the minimum and maximum % bound was also obtained for the fit of each binding isotherm.

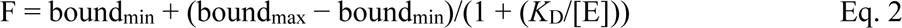

### Processivity assay

Annealed 8 bp duplexes were added to assay buffer containing 25 mM HEPES pH 7.5, 10 mM MgCl_2_, 3 mM DTT, 25 mM NaCl and 0.2 mg/ml BSA and the indicated NTP at a concentration of 0.5 mM. The reactions were initiated by adding 1μl Thg1/TLP enzyme (to a final concentration of 15 μM) and were incubated at room temperature to initiate 5’-extension. A cold chase step was performed by adding 5 μM unlabeled RNA substrate competitor at indicated time points. The reactions were quenched after allowing the reactions to continue for desired time points by adding 100 mM EDTA to a final concentration of 10 mM. The products were then resuspended in formamide loading dye and 42.8% DMSO followed by heat denaturation at 95 °C for 5 min. Reactions were resolved, visualized, and quantified as described above.

## Results

### Simple RNA duplexes are substrates for 3’-5’ addition activity of Thg1/TLPs

Although purified Thg1/TLP enzymes have been assayed using a variety of *in vitro* activity assays, these assays nearly all utilized highly structured tRNA molecules or tRNA-like model substrates. Moreover, consistent comparison of catalytic preferences across the diverse landscape of Thg1/TLP enzymes has been difficult due to differences in sequence, structure and site of nucleotide addition with the tRNA or tRNA-like substrates tested in assays performed to date (Rao et al. 2011; Smith and Jackman 2012; Desai et al. 2018; Patel et al. 2021). Thus, we sought to develop a standardized assay using a simplified RNA substrate for 3’-5’ addition as a common platform for evaluating enzyme activities across Thg1/TLP family members. We also sought to create a system that would be amenable to analysis of 3’-5’ addition using gel electrophoresis, as opposed to many assays used so far for 3’-5’ addition, which typically rely on thin-layer chromatography to detect 5’-end oligonucleotide fragments generated by nuclease digestion of labeled tRNA substrates. Here, we tested whether a relatively short RNA could be utilized as a substrate to investigate 3’-5’ addition reactions by annealing with a separate oligonucleotide to provide the template sequence for 3’-5’ addition, similar to the role of the 3’-end of the tRNA that serves as a template for 3’-addition in the context of mitochondrial tRNA 5’-editing (**Figure 1**).

The substrate RNA chosen for initial assays was a 26-mer derived from the sequence following the T7 promoter in a pET plasmid (see Methods). This RNA has a relatively balanced nucleotide composition (19% U, 23% A, 27% G, 31% C) and is not predicted to form stable secondary structures at 27 °C. For these assays, the RNA 26-mer substrate was uniformly labeled by *in vitro* transcription in the presence of [α-^32^P] ATP. Importantly, the *in vitro* transcribed RNA contains a 5’-triphosphorylated end, which provides a pre-existing 5’-leaving group for 3’-5’ nucleotide addition (**Figure 2A**). Thus these assays do not require prior activation of the RNA substrate by 5’-adenylyation as is required for Thg1/TLP activity on 5’-monophosphorylated RNA, and thus allowed us to leave ATP out of the reactions (Jackman and Phizicky 2008; Smith and Jackman 2012). The labeled 26-mer RNA substrate was annealed to an RNA template oligonucleotide complementary to first 19 nucleotides of the model RNA substrate, creating an 8-nucleotide (8-nt) overhang with alternating GC nucleotides that could serve as a template for 3’-5’ polymerase activity (**Figure 2A**). Prior to annealing with 26-mer substrate, the *in vitro* transcribed RNA template was treated with phosphatase to generate a 5’-hydroxyl end, thus preventing any potentially competing extension of the 5’-end of the unlabeled RNA template during the reactions.

**Figure 2:**
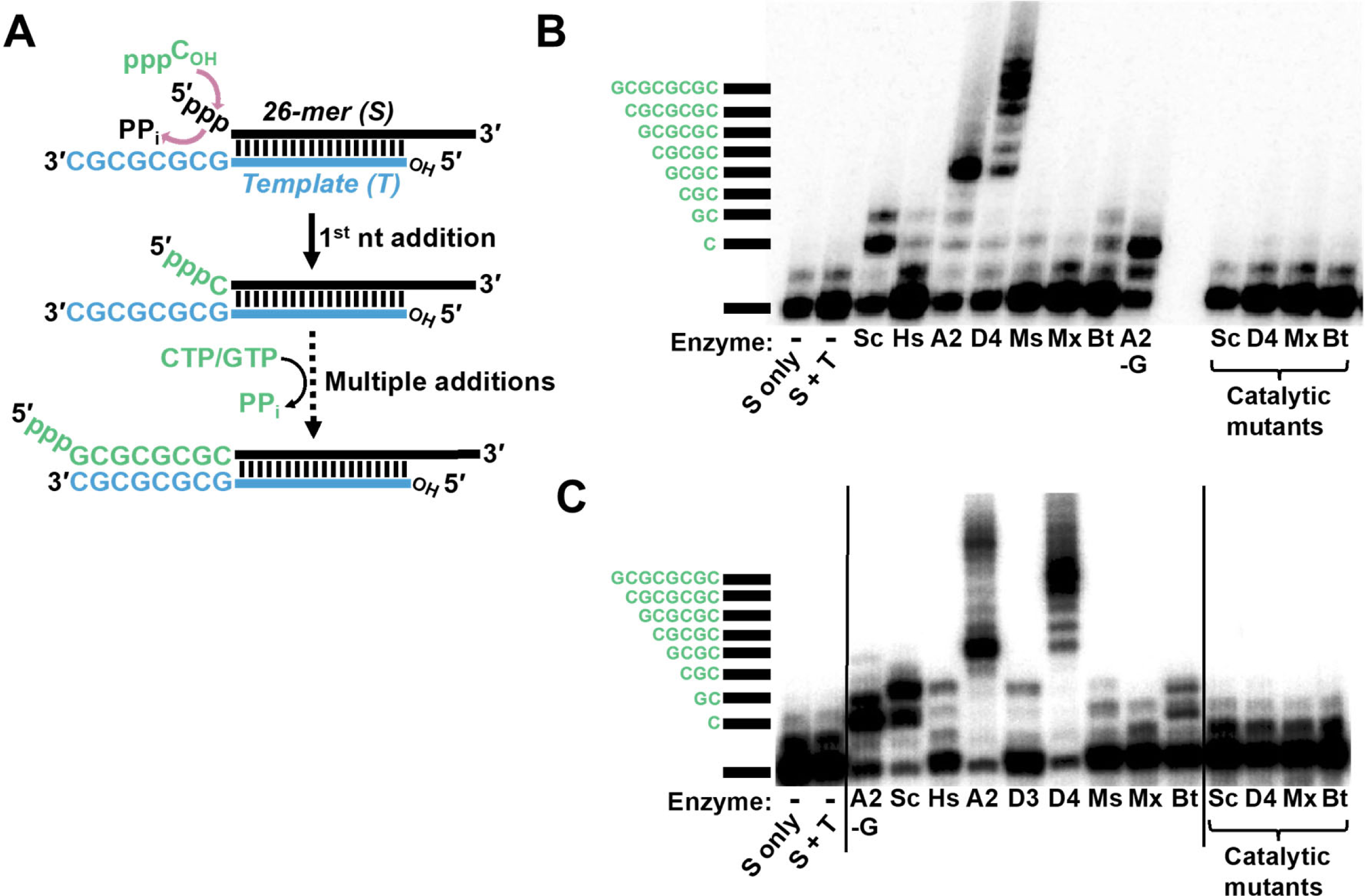
Targeted 3’-5’ polymerase activity on RNA duplexes catalyzed by Thg1/TLP enzymes. **(A)** Schematic of the reaction catalyzed with RNA duplexes, in which a radioactively labeled 5’-triphosphorylated 26-mer RNA substrate transcript (S) is annealed to a 5’-hydroxylated (unlabeled) RNA template oligonucleotide (T). The annealed duplexes contain an 8-nucleotide 3’-overhang (5’-GCGCGCGC-3’) to serve as a template for 3’-5’ polymerase activity of the tested Thg1/TLP enzymes. Template-dependent 3’-5’ addition of either CTP or GTP (depending on the templating nucleotide as shown) will result in incorporation of nucleotides into the 5’-end of the labeled S, and these elongated products are visualized by denaturing gel electrophoresis. **(B)** A survey of purified Thg1/TLP enzyme activities with the 19 bp RNA duplex, as shown in (A). Reactions were performed as described in Methods, with individual enzymes indicated under each lane (Sc: ScThg1, Hs: HsThg1, A2: AcaTLP2, D4: DdiTLP4, Ms: MsTLP, Mx: MxTLP and Bt: BtTLP). The A2-G lane corresponds to a reaction where GTP has been omitted and CTP is the only nucleotide added to the reaction. The last four lanes contain purified catalytically inactive Thg1/TLP variants, as indicated. The first two lanes of the gel are no enzyme controls indicating the position of the unreacted 26-mer substrate (lane S), and of the unreacted duplex (lane S + T), which demonstrates that migration of the substrate is not affected by presence of the template RNA on this denaturing (20% polyacrylamide/8 M urea) gel. The thick black line marks the position of the unreacted substrate, and bands corresponding to nucleotide addition products are labeled accordingly. **(C)** A survey of purified Thg1/TLP enzyme activities with the 8 bp RNA duplex, performed as described in Methods. Individual enzymes were indicated as described above, with the addition of D3: DdiTLP3 for this assay.

First, we tested the ability of seven different purified Thg1/TLP enzymes to catalyze templated 5’-extension of this duplex substrate. Enzymes were chosen to represent previously characterized enzymes from all three domains of life that are readily expressed and purified from *E. coli* and have been characterized in other *in vitro* assays. These included eukaryotic enzymes such as *S. cerevisiae* Thg1 (ScThg1), *Homo sapiens* Thg1 (HsThg1), *A. castellanii* TLP2 (AcaTLP2) and *D. discoideum* TLP4 (DdiTLP4), as well as the archaeal TLP from *Methanobrevibacter smithii* TLP (MsTLP), and bacterial TLPs from *Myxococcus xanthus* (MxTLP) and *Bacillus thuringiensis* (BtTLP). Annealed RNA duplexes were incubated with each enzyme in the presence of CTP and GTP, according to the GC template sequence, and analyzed for addition of 5’-nucleotides by resolving the reaction products on denaturing polyacrylamide gels (**Figure 2B**).

Figure 2B shows the results of an optimized assay capable of resolving products of 3’-5’ addition to single nucleotide resolution, which necessitated addition of 42.8% DMSO and equimolar amounts of unlabeled 26-mer RNA substrate (to compete away template from the labeled RNA). These strategies have both been used to improve resolution in other assays of RNA-dependent RNA polymerases that generate highly stable RNA:RNA duplexes (Arnold et al. 1999; Nwokeoji et al. 2017). However, identifying the precise number of nucleotides added by each enzyme remained a challenge with a standard RNA transcript because of heterogeneity observed with the *in vitro* transcript derived from run-off transcription, as has been previously described (Gholamalipour et al. 2018) (**Figure S1A**). Thus, we engineered a clone with an HDV ribozyme sequence incorporated immediately 3’-to the RNA substrate, which enabled transcription of a nearly homogeneous product (∼95% major band) to be used in the assay, thus enabling robust resolution of multiple addition products in the activity assays (**Figure S1B**).

Using the optimized RNA substrate, we observed varying degrees of enzyme-dependent products that migrated higher than the unreacted substrate in reactions with all tested enzymes (Figure 2B). However, a clear pattern of longer products suggesting multiple nucleotide incorporation was observed with two TLPs, AcaTLP2 and DdiTLP4. Importantly, the longer extension products generated by AcaTLP2 were not observed in reactions performed under conditions where GTP was omitted from the reactions and only CTP was provided (Figure 2B, compare lane A2-G with lane A2), indicating that templated incorporation of both GTP and CTP was required to observe robust 5’-end extension by this enzyme. The same results were observed for reactions with DdiTLP4 that contained only CTP (data not shown). The reaction with GTP-omitted also allowed us to identify the band corresponding to the first CTP addition and thus to individually identify bands corresponding to each subsequent addition (up to and beyond the expected 8-nt 5’-additions to the substrate RNA). Interestingly, we consistently saw distinct patterns of activity for DdiTLP4 vs. AcaTLP2. DdiTLP4 appears to substantially terminate addition at or near the end of the template sequence, based on the majority of the extended products corresponding to the band with 8-9 nt added to the 5’-end of the substrate. AcaTLP2 on the other hand exhibits a prominent stop earlier in the reaction (after 4 nt additions), although minor bands corresponding to even longer products can be observed in these assays.

We verified that all of the enzyme-dependent products observed on the gels were indeed due to the 3’-5’ polymerase activity of each enzyme using catalytic mutants of ScThg1, DdiTLP4, MxTLP and BtTLP in which an essential metal-binding carboxylate residue (D76 in ScThg1, or its analogous residue in the other enzymes) is mutated to alanine, rendering the corresponding proteins completely inactive (Jackman and Phizicky 2008; Hyde et al. 2010; Smith and Jackman 2012). The results with catalytically inactive mutants were identical to the no enzyme control reactions (Figure 2B), confirming that the reaction products observed with the active Thg1 and TLP enzymes are enzyme-dependent 3’-5’ addition products. Thus, we conclude that the RNA duplex with a base-paired length of 19 nucleotides can undergo 5’-extension of the 26-mer model RNA in a template-directed manner, and that two enzymes (DdiTLP4 and AcaTLP2) appear to catalyze the reaction with the highest efficiency.

### TLPs accommodate RNA duplexes of varying length for 5’-targeted extension

Shorter RNA duplexes could provide several advantages for the use of 3’-5’ polymerases to incorporate 5’-nucleotides in this template-directed approach. First, disruption of RNA:RNA duplex reaction products by electrophoresis is likely to be more favorable with fewer complementary base pairs. Second, the use of shorter RNA template oligonucleotides will provide advantages in terms of cost and possible ease of chemical synthesis. Thus, we sought to test whether shorter duplexes could be accommodated by Thg1/TLP enzymes. Five RNA templates were designed to test duplex lengths from 7-19 base pairs by varying the start site for *in vitro* transcription for each construct (Figure 3A). The resulting templates all base-pair with the 26-mer RNA substrate to create the same 8-nt overhang (GCGCGCGC) for nucleotide incorporation. Template-dependent 5’-extension was performed for the five RNA duplexes with AcaTLP2 and DdiTLP4, as well as with ScThg1, as an example of an enzyme that appeared to catalyze less efficient 3’-5’ extension (Figure 3B). Interestingly, we observed that duplexes with as few as 7 base pairs between the RNA substrate and RNA template were utilized even more efficiently than the longer RNA/RNA duplexes tested initially. Again, DdiTLP4 extended each substrate with a major product that corresponds to the expected 8-nt addition, although the overall efficiency appeared to improve with shorter duplexes. For AcaTLP2, likewise, the major 4-nt 5’-extension product was observed with all duplexes, but with the shorter duplexes, AcaTLP2 was able to clearly extend further to create longer products than DdiTLP4 (∼10-12 nt extended).

**Figure 3:**
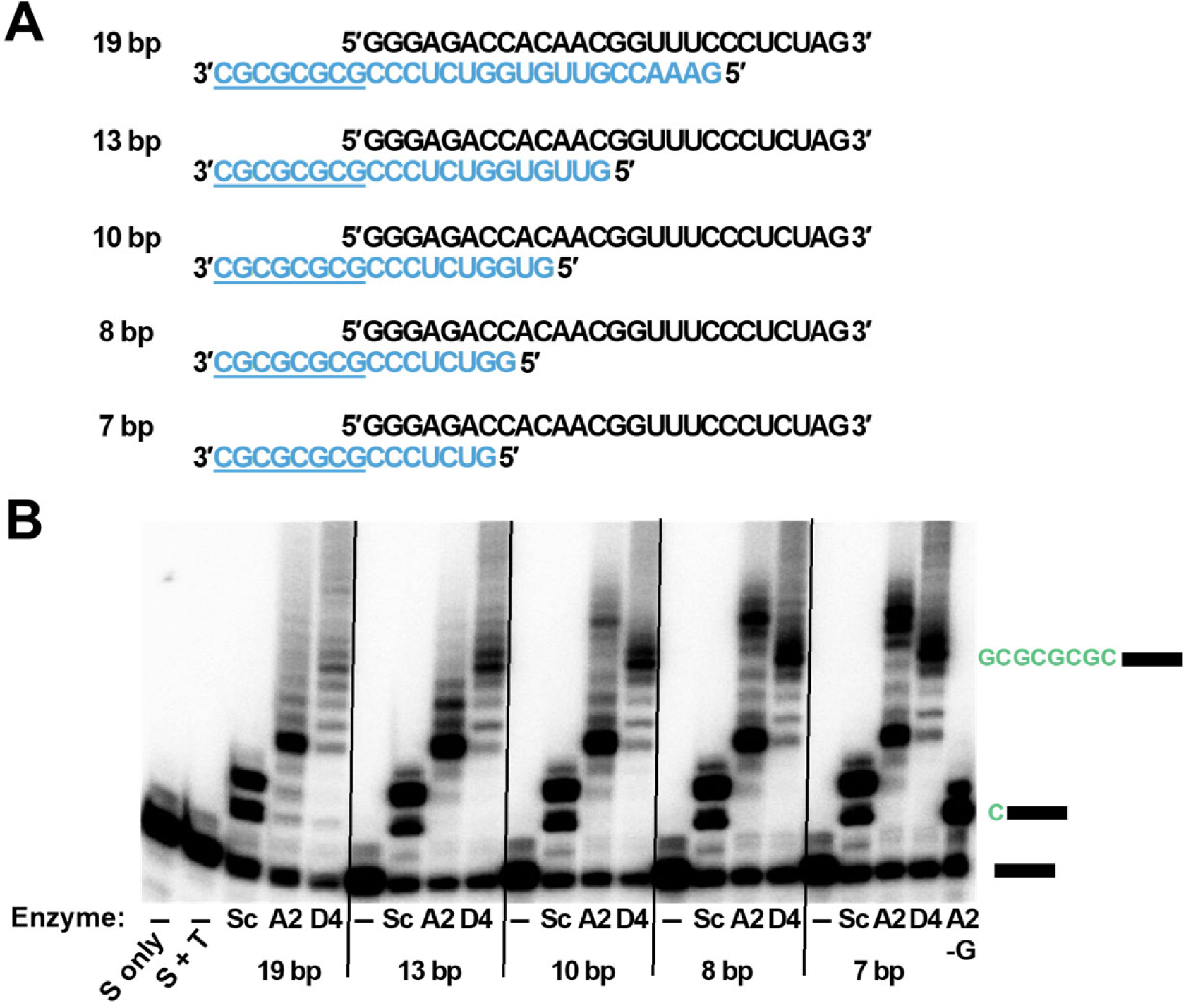
AcaTLP2 and DdiTLP4 catalyze 3’-5’ polymerase activity with RNA duplexes of varying base paired length. **(A)** Design of duplex RNAs using the same 26-mer RNA substrate (shown in black) and template sequences of varied length (shown in blue) to generate varied base paired regions of 7-19 bp (highlighted in pink) **(B)** Assays with the indicated duplexes from 19 to 7 bp are shown using three different Thg1/TLP enzymes (Sc: ScThg1, A2: AcaTLP2, D4 (DdiTLP4). The band corresponding to single C-addition product is shown in the last lane (A2-G), and migration of the unreacted substrate only (S only) and substrate in the presence of template (S+T) are also shown, as described in Figure 2.

This result prompted us to repeat the earlier survey of activities with the full panel of Thg1/TLP enzymes (Figure 2C). Although the overall trends of activity were the same for each enzyme comparing 19 bp (Figure 2B) vs. 8 bp (Figure 2C) duplex substrates, all of the tested enzymes exhibited slightly more robust nucleotide addition with the shorter duplexes. Interestingly, the longer addition products were again observed with AcaTLP2 that seem to extend beyond the full-length (8-nt) addition products. Whether these additional nucleotides added by AcaTLP2 are the result of templated or non-templated 3’-5’ addition cannot be determined by this assay. For ScThg1, HsThg1, DdiTLP3, MsTLP, MxTLP and BtTLP, the major products that were observed run at or near the position of the first CTP addition, indicating that each of these enzymes incorporates only 1-2 nucleotides with this RNA duplex. Moreover, the slight differences in mobility of the single nucleotide addition products observed with some of these less efficient enzymes may suggest that some amount of non-Watson-Crick incorporation could be occurring in these reactions. Catalytically inactive mutants of each of the tested enzymes were unable to catalyze 5’-extension, as expected.

### Efficient 5’-extension correlates with high affinity binding to RNA duplexes

To determine whether some of the differences in Thg1/TLP activity observed in the extension assays could be due to differences in binding affinity for these RNA duplexes, we used a double filter binding assay to determine *K_D_* values for the set of five RNA duplexes of varied length. To compare the observed binding affinities with those for tRNA substrates, we also determined *K_D_* of each enzyme for a bacterial tRNA^His^ transcript from *M. xanthus* (Mx-tRNA^His^). Thg1/TLP enzymes generally bind to tRNA with *K_D_* values in the μM range, as observed using both biochemical and kinetic methods (Smith and Jackman 2012; Patel et al. 2021). As a control, ScThg1, AcaTLP2, DdiTLP4, MsTLP and BtTLP were first tested for binding to the single-stranded 26-mer RNA substrate in the absence of an RNA template (Figure 4A). For DdiTLP4, BtTLP and MsTLP, no detectable binding was observed to the model RNA substrate alone, even at concentrations as high as 10 μM in the assays. AcaTLP2 and ScThg1 both exhibited some non-specific binding to the 26-mer RNA alone, as evident from the linear increase in bound RNA (the concentration of enzyme was at least 10^4^-fold greater than the labeled duplex in these assays). Overall, these results suggest that Thg1/TLP enzymes require more complex structural features to interact stably with RNA and do not bind efficiently to single-stranded RNA, which is consistent with all known biological substrates for the enzyme identified to date.

**Figure 4:**
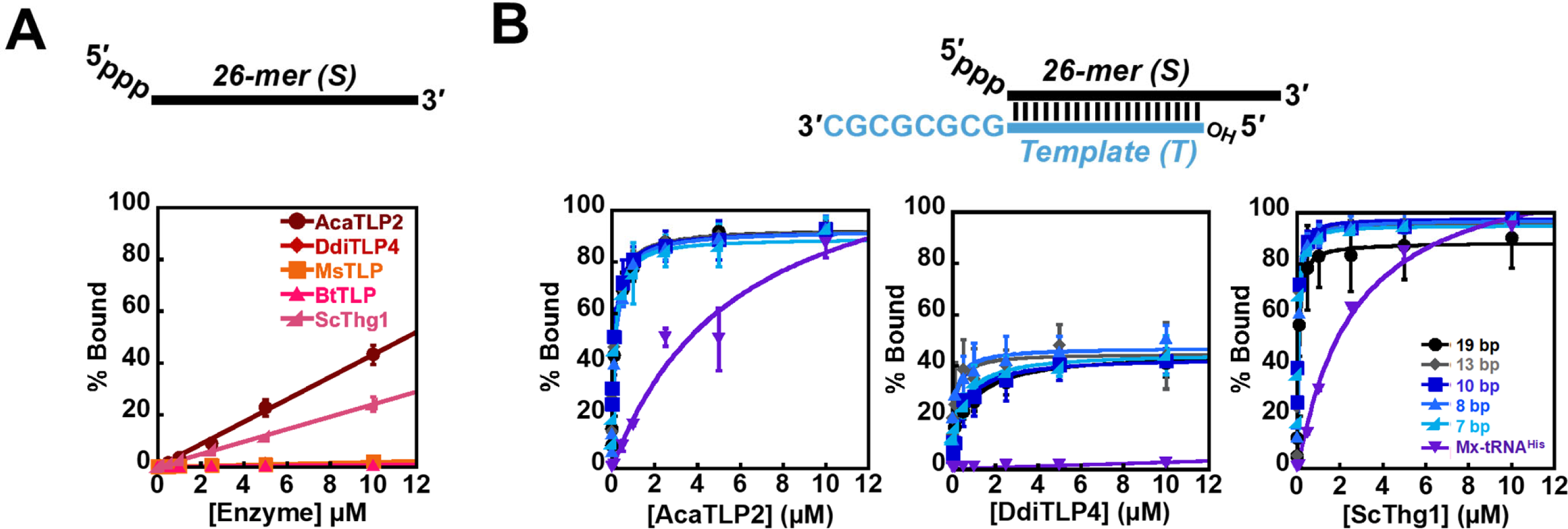
Thg1/TLP enzymes bind RNA duplexes with high affinity. **(A)** Binding to the single-stranded model RNA substrate (26-mer) was measured using a double filter-binding assay, with the indicated enzymes (AcaTLP2, DdiTLP4, MsTLP, BtTLP, and ScThg1) representing diverse Thg1/TLPs from Eukarya, Archaea and Bacteria. The fraction of bound RNA product was plotted vs. the concentration added in each reaction (0-10 μM), with linear fits representing non-specific binding (*K_D_* > 10μM) for enzymes that exhibited detectable RNA binding in the assays. Average fraction bound was determined from triplicate assays performed with each enzyme. **(B)** Binding to RNA duplexes with varied base-paired length (from 7-19 bp) and *M. xanthus* tRNA^His^ was measured using a double filter-binding assay. Average fraction bound was determined from triplicate assays performed with AcaTLP2, DdiTLP4 and ScThg1, as indicated, and plotted vs. concentration of each enzyme added (0-10 μM). The data were fit to Eq. 2 to determine the apparent *K_D_* for each enzyme binding to the tested RNAs, which are indicated in Table 1.

Binding was then measured using the same assay for each of the annealed RNA duplexes and compared to Mx-tRNA^His^ (Figure 4B). The varied length of the RNA duplex between 7-19 base pairs did not significantly affect the overall affinity observed for any tested enzyme (**Table 1**). Therefore, the increase in apparent catalytic efficiency observed with shorter RNA duplexes (Figure 3) is not due to increased binding affinity. Moreover, DdiTLP4 and AcaTLP2, which both exhibited the most robust 3’-5’ polymerase activity with duplex substrates also bind to these RNAs with notably higher affinity than the other weakly active TLPs (BtTLP and MsTLP) (**Table 1** and **Figure S2**). Strikingly, the affinity of AcaTLP2 for the duplex RNAs were substantially higher than any tested Thg1/TLP’s affinity for tRNA, with *K_D_* values between 160-210 nM for all five RNA duplexes, in contrast to the typical μM binding constants that have been observed for Thg1/TLP enzymes binding to tRNA, and also seen here for Mx-tRNA^His^ (**Table 1**). The other striking result from this experiment was the unexpectedly low *K_D_* values (<100 nM) exhibited by ScThg1 for the RNA duplexes, binding even more tightly than the highly active DdiTLP4 and AcaTLP2 to these RNAs (Figure 4 and **Table 1**). Since ScThg1 exhibits very poor 5’-extension activity with the RNA duplexes (Figure 2B), the catalytic relevance of the tight-binding complexes between ScThg1 and the RNA duplexes (tighter even than the ScThg1 interaction with its physiologically relevant tRNA^His^ substrate) is not yet clear.

**Table 1:** *K_D_* values for Thg1/TLP binding to RNA duplexes measured by filter-binding assay.

| RNA | AcaTLP2<sup>a</sup><br>(μM) | DdiTLP4<sup>a</sup><br>(μM) | MsTLP<sup>a</sup><br>(μM) | BtTLP<sup>a</sup><br>(μM) | ScThg1<sup>b</sup><br>(μM) |
| --- | --- | --- | --- | --- | --- |
| 19 bp | $0.17 \pm 0.02$ | $1.0 \pm 0.3$ | $8.0 \pm 2.4$ | $6.8 \pm 1.5$ | $0.065 \pm 0.008$ |
| 13 bp | $0.18 \pm 0.04$ | $0.2 \pm 0.1$ | $3.9 \pm 0.8$ | $12.1 \pm 4.1$ | $0.042 \pm 0.002$ |
| 10 bp | $0.21 \pm 0.05$ | $0.6 \pm 0.2$ | $2.1 \pm 0.3$ | $13.5 \pm 2.2$ | $0.061 \pm 0.008$ |
| 8 bp | $0.19 \pm 0.03$ | $0.2 \pm 0.1$ | $3.5 \pm 0.6$ | $9.7 \pm 1.2$ | $0.075 \pm 0.008$ |
| 7 bp | $0.16 \pm 0.03$ | $0.7 \pm 0.1$ | $3.7 \pm 0.8$ | $11.5 \pm 4.3$ | $0.054 \pm 0.010$ |
| Mx-tRNA<sup>His</sup> | $5.88 \pm 3.12$ | $>10^c$ | $1.7 \pm 0.2$ | $>10^c$ | $2.6 \pm 0.5$ |

<sup>a</sup>Average fraction bound at each enzyme concentration was calculated from three independent experiments, plotted vs. enzyme concentration to calculate $K_D$ using eq.1.
<sup>b</sup> $K_D$ values for ScThg1 were calculated using only two independent experiments.
<sup>c</sup>A lower limit to $K_D$ was estimated for DdiTLP4 and BtTLP based on the highest enzyme concentration used in the assays.

### Kinetic analysis of AcaTLP2 and DdiTLP4 reveals efficient 3’-5’ polymerase activity with RNA duplexes

To further evaluate the catalytic proficiency of TLPs in comparison to other activities catalyzed by the Thg1/TLP family, we measured the single turnover observed rates with the two highly active TLPs and the two shortest RNA duplexes (8 bp and 7 bp) that were most efficient substrates for these enzymes (Figure 5). These assays were performed with at least 100-fold excess enzyme over each RNA duplex, with saturating (0.5 mM) GTP and CTP (Figure 5A). For each time point, the total amount of product formation was quantified by summing all 3’-5’ addition products created by each enzyme and plotting this fraction product vs. time to yield the *k_obs_* (min^-1^) for each enzyme (Figure 5B). Calculated observed rates (*k_obs_*) show that DdiTLP4 is modestly more proficient than AcaTLP2 for addition to both 7 bp and 8 bp RNA duplexes (Figure 5 **and Table 2)**. Consistent with the end point assays conducted earlier (Figure 3), neither enzyme exhibited a significant difference in observed rate between the 7 bp vs. 8 bp duplexes (**Table 2**). The observed rates were also compared to the previously measured rates of multiple nucleotide addition to a 5’-truncated tRNA by AcaTLP2 and DdiTLP4 (Long and Jackman 2015). Based on this comparison, AcaTLP2 shows a stronger catalytic preference for repairing 5’-truncated tRNA than for adding multiple nucleotides to the RNA duplexes tested here. DdiTLP4 however, exhibits faster rates with the RNA duplexes over 5’-truncated tRNA.

**Figure 5:**
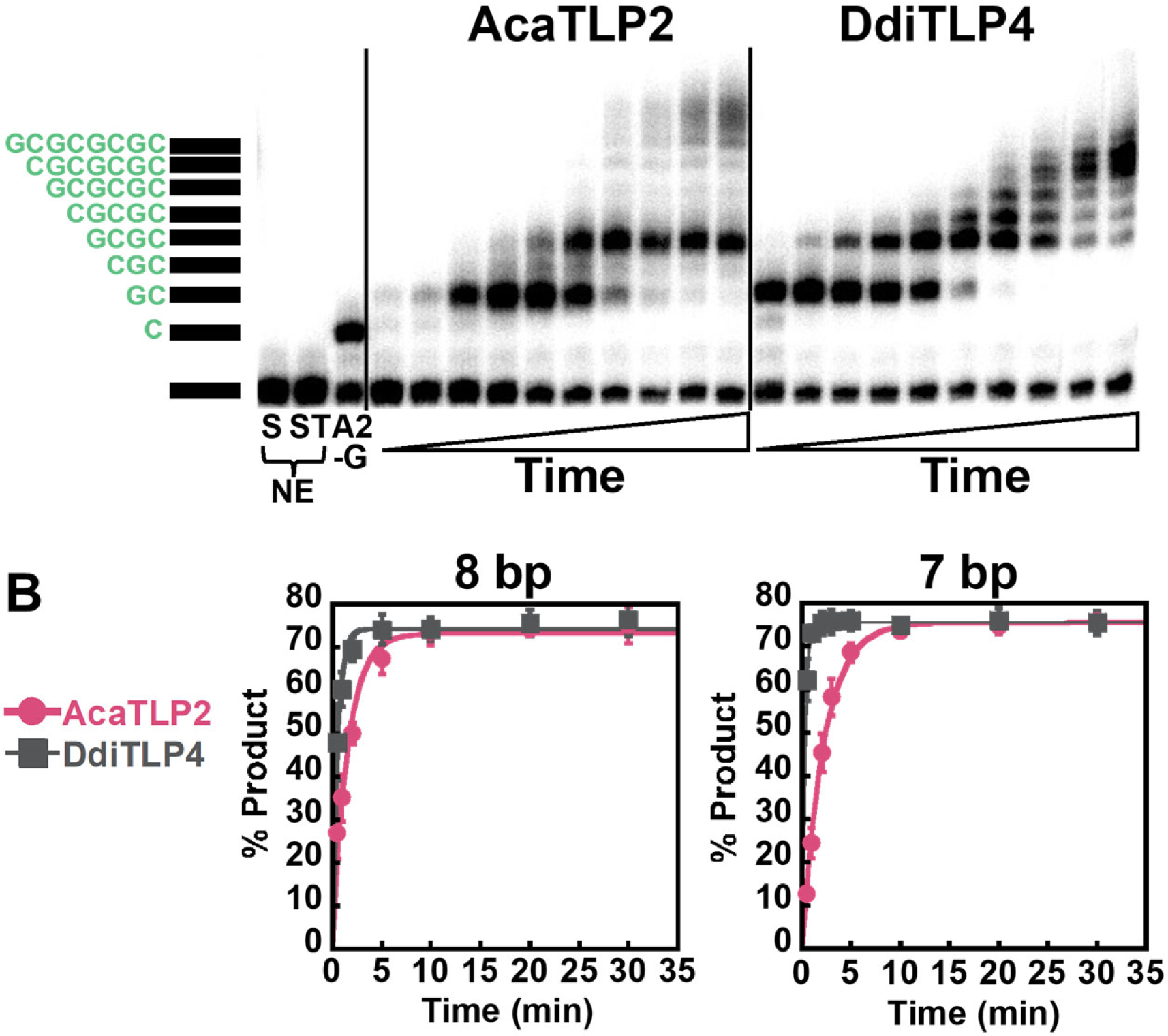
TLPs catalyze efficient 3’-5’ polymerase activity with RNA duplexes. **A)** A representative time course of 3’-5 polymerase activity measured with AcaTLP2 and DdiTLP4 (15 μM) with the 7 bp RNA duplex assayed under single turnover conditions (at least 100-fold excess [Enzyme] over [RNA]) with individual time points measured from 0-120 minutes (time points shown in this assay correspond to 0.5, 1, 2, 5, 10, 20, 30, 45, 60, 120 min). The band corresponding to unreacted substrate (S) or substrate in reactions that contain template (ST) is indicated by the black bar on the side of the time course. Extension products are as indicated by added nucleotides shown in green. Reaction A2-G shows the migration of the single nucleotide addition product observed in the absence of GTP, as shown previously. **(B)** Determination of *k_obs_* values for 15 μM enzyme (AcaTLP2, circles; DdiTLP4, squares) from the average percent total nucleotide addition products observed in triplicate reactions measured with the 8 bp and 7 bp substrates, as indicated. The data were fit to Eq. 1 to yield *k_obs_* for each enzyme with the indicated RNA.

**Table 2:** Single turnover multiple nucleotide incorporation rates *(k_obs_*) for Thg1/TLP activity.

| Enzyme | 8 bp<sup>a</sup><br>(min<sup>-1</sup>) | 7 bp<sup>a</sup><br>(min<sup>-1</sup>) | 5'-truncated tRNA<br>(min<sup>-1</sup>) | Full-length tRNA<sup>His</sup>A<sub>73</sub><br>(min<sup>-1</sup>) |
| --- | --- | --- | --- | --- |
| ScThg1 | ND<sup>b</sup> | ND | ND | 3.6 ± 0.3 |
| AcaTLP2 | 0.63 ± 0.06 | 0.45 ± 0.02 | 4.9 ± 2.2 | NT<sup>c</sup> |
| DdiTLP4 | 1.9 ± 0.1 | 3.45 ± 0.06 | 0.71 ± 0.05 | NT<sup>c</sup> |

<sup>a</sup> Average percent product conversion at each time point was calculated from three independent experiments, plotted vs. time and fit to a single exponential rate equation to calculate $k_{obs}$ using eq.2
<sup>b</sup> ND = no activity detected for this substrate
<sup>c</sup> NT = not tested

In the time courses, we noted a distinct pattern of nucleotide addition products in which some reaction intermediates accumulated prominently at early time points and appear to be subsequently converted into a longer product (Figure 5A). The apparent production of these prominent intermediates could reflect inherent differences in reactivity for certain positions in the elongating substrate:template duplex, or a propensity for the enzyme to exhibit a dissociative type behavior that leads to release of incompletely extended products during the reactions. Although multiple aspects of the overall mechanism and kinetic behavior of several 3’-5’ polymerases have been studied extensively, a kinetic assessment of enzyme processivity (i.e., how many nucleotides are added in one enzyme-substrate encounter) has not been carried out previously for any Thg1/TLP enzyme.

To test whether AcaTLP2 and DdiTLP4 exhibit distributive kinetic behavior in extending RNA duplexes, we carried out an assay designed to reveal whether the enzyme is undergoing multiple cycles of release and rebinding of substrates, limiting the production of longer reaction products in some assays (Figure 6). Reactions were initiated with addition of enzyme as before, but at certain times after initiation, reactions were chased by the addition of 5 μM unlabeled RNA 26-mer substrate. Reactions were allowed to continue for a total of either 5 min or 15 min, corresponding to time points where the intermediate products (2-nt and 4-nt addition) are robustly observed (Figure 5). Control reactions were performed with no addition of unlabeled substrate, and analysis of the 5 min reactions showed primarily 2-nt product produced by AcaTLP2 and a mixture of both 2-nt and 4-nt product by DdiTLP4, consistent with the products observed for the same time point in Figure 5 (4^th^ time point in each time course). Likewise, the unchased reactions allowed to proceed for 15 min showed the expected pattern of product formation. Measurement of the extent of product formation in reactions that contained added unlabeled competitor RNA can reveal whether the 3’-5’ polymerase remained bound to the same labeled substrate:template duplex during the time course, or whether the enzyme may have released and re-bound the duplex, which is more likely to be an unlabeled RNA after the chase. A fully processive enzyme would yield the same amount of labeled product formation regardless of the presence or timing of addition of the competing unlabeled substrate RNA. However, a decrease in full-length product formation in the reactions where unlabeled competitor was added compared to the control could reveal that the enzyme is acting more distributively.

**Figure 6:**
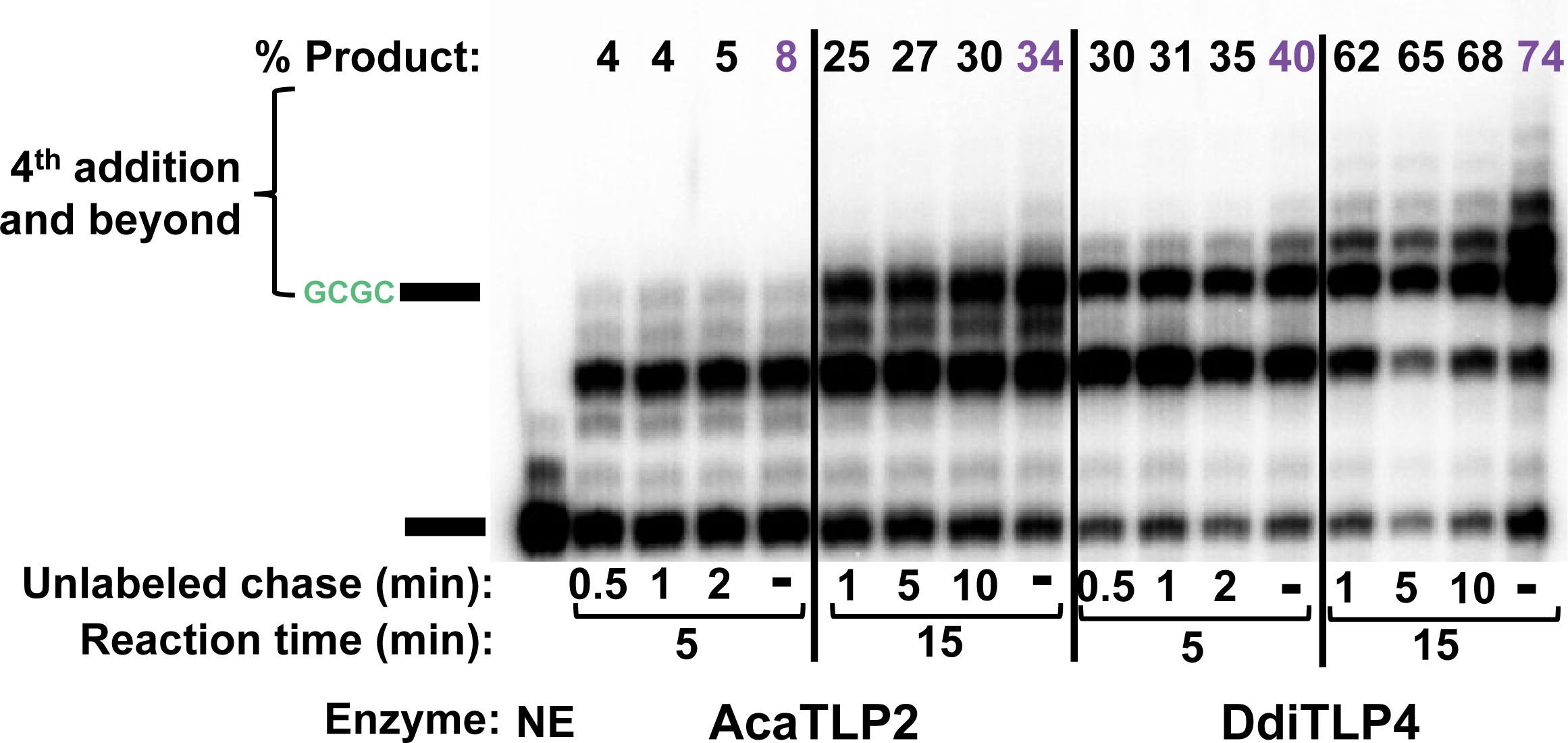
Assessment of processivity exhibited by AcaTLP2 and DdiTLP4 during reaction with RNA duplex substrates. Processivity assays using the 8 bp duplex substrates were performed as described in Methods. Briefly, reactions were initiated by addition of 15 μM of the indicated enzyme and with duplexes containing only labeled 26-mer substrate RNA. At the times indicated for the unlabeled chase (0.5-10 min after each reaction was initiated) 5 μM of the unlabeled RNA duplex was added to each reaction. Reactions indicated by – correspond to no unlabeled RNA chase added. Reactions were allowed to proceed for a total of either 5 or 15 minutes, as indicated below each panel, at which point they were quenched and resolved as described above. The percent of addition products longer than the 4 nt-extended intermediate were quantified for each reaction and are indicated above each lane.

For each reaction with added unlabeled RNA, the total percent of products formed beyond the accumulated 4-nt intermediate was quantified and compared with the relevant control reaction. For each set of reactions, we did observe a slightly greater amount of longer extension products in the control (no unlabeled chase) compared to reactions that had been chased with unlabeled RNA. The effects were most pronounced in the 15 min reaction products, with a significantly higher percent of longer products in the control reactions than in each of the corresponding chased reactions. Although these results may indicate a modest amount of distributive behavior exhibited by AcaTLP2 and DdiTLP4 during duplex extension, the results observed here do not support a distributive mechanism as the main driver for accumulation of specific intermediates during the course of the 3’-5’ polymerase reactions with these duplex substrates.

## Discussion

Here we demonstrate that small RNA duplexes can serve as substrates for 3’-5’ polymerases of the Thg1/TLP superfamily. Consistent with previous *in vitro* studies that suggested relatively more flexibility of TLPs in terms of substrate recognition and multiple nucleotide incorporation into tRNAs compared to Thg1, two TLPs (AcaTLP2 and DdiTLP4) were the most proficient at extending the 5’-end of the 26-mer RNA substrate when duplexed to a templating RNA oligonucleotide (Figure 2). Both enzymes were capable of fully extending the RNA substrate utilizing the provided alternating G/C nucleotide templates to direct CTP and GTP incorporation. Moreover, AcaTLP2 and DdiTLP4 catalyze 3’-5’ polymerase activity with an RNA duplex containing as few as 7 base pairs of overlap with the RNA substrate, and in fact exhibit even greater extension efficiencies with these substrates than for longer duplexes containing base paired regions of up to 19 bp (Figure 3, **Table 1** and **Table 2**). These assays establish 3’-5’ polymerases as a potential versatile tool for broader post-transcriptional nucleic acid 5’-end labeling reactions by virtue of their ability to act on these highly simplified substrates, suggesting that the 3’-5’ polymerase activity of TLPs could have important biotechnology applications.

These studies are not the first attempt to exploit the 3’-5’ extension activity of TLPs to add nucleotides to engineered RNA substrates. Previously, a split-tRNA approach has been successfully used to guanylylate an RNA substrate of interest, where the tRNA is divided at the D-loop and a guide RNA oligonucleotide is provided to make up for an overall tRNA structure appropriate for TLP recognition (Desai et al. 2018). Additionally, an alternative approach successfully utilized a guide RNA oligonucleotide to disrupt the tRNA structure in the acceptor stem and the D-loop region, thus providing an alternative template sequence to guide multiple nucleotide additions to a standard tRNA substrate (Chen et al. 2019). While these experiments have pointed out the fact that TLPs are amenable to be engineered as post-transcriptional RNA 5’-end labeling tools, these studies are the first, to our knowledge, to completely divorce 3’-5’ polymerase chemistry from something resembling aspects of tRNA structure.

The *bona fide* biological substrates for the enzymes that act most proficiently on these RNA duplexes are not yet known. Interestingly, however, the observed *in vitro* preferences for AcaTLP2 and DdiTLP4 to bind to the RNA duplexes over binding to tRNA (Figure 4 and **Table 1**), combined with the proficient kinetic behavior of these enzymes with the tested duplexes that is similar to (AcaTLP2), or even exceeds (DdiTLP4) the observed rates with tRNAs (Figure 5 and **Table 2**) may reflect an ability of these enzymes to act on non-tRNA substrates *in vivo*. The strong binding affinity exhibited by both enzymes for RNA duplexes may also reflect some structural feature of a biologically relevant substrate that has not yet been conclusively identified for either enzyme.

These data suggest that AcaTLP2 and DdiTLP4 are strong candidates for engineering TLPs to utilize 3’-5’ polymerase activity in a targeted manner to incorporate desired 5’-end sequences into specific RNAs by annealing a templating RNA oligonucleotide (possibly to as few as 7 bp) to the RNA 5’-end. A few interesting possibilities for exploiting reverse polymerization in the future could include post-transcriptional site-specific 5’-end labeling, incorporating labeled/modified nucleotides or introducing affinity/fluorescence or localization tags in a single reaction. Along these lines, we have already been able to use 3’-5’ polymerases to add a single non-GTP nucleotide to the 5’-end of RNAs post-transcriptionally, which has been a known challenge associated with T7 RNA polymerase due to its inherent preference to start transcription with a G nucleotide (Smith and Jackman 2014; Mullins et al. 2023).

The high affinity (low nM) interaction of ScThg1 with RNA duplexes compared to its affinity for tRNA^His^ was unexpected (Figure 4 and **Table 1**). It is well-established that ScThg1 exhibits a strong selection for activity on tRNA^His^ over other tRNA substrates, consistent with the biological importance of selectively adding the G_-1_ identity element for HisRS to this tRNA, in order to ensure accurate aminoacylation of tRNA^His^ (Cooley et al. 1982; Rudinger et al. 1997; Rosen and Musier-Forsyth 2004; Jackman and Phizicky 2006a). It is intriguing that ScThg1 exhibited the highest affinity for the short RNA duplexes of all tested enzymes despite its inability to incorporate multiple nucleotides into these RNAs (Figure 2, Figure 4 and **Table 1**). Although the binding of ScThg1 to the RNA duplexes clearly does not result in formation of a catalytically productive complex for multiple nucleotide incorporation, the molecular basis for this strong specific interaction between ScThg1 and RNA duplexes remains to be determined.

The archaeal and bacterial TLP enzymes tested here (MsTLP and BtTLP) exhibited weak activity and poor extension beyond 1-2 nucleotides with the RNA duplex substrates (Figure 2), revealing that the ability to act on these relatively simple duplexes is not a common property of all members of the TLP family. Interestingly, however, BtTLP efficiently repairs 5’-truncated tRNAs with catalytic efficiency similar to AcaTLP2 and DdiTLP4, indicating that trends of substrate specificity are unique to each enzyme (Rao et al. 2011). Previously, differences in the extent of nucleotide incorporation by different TLPs (from *M. xanthus*, *Methanothermobacter thermoautotrophicus*, *Methanosarcina barkeri*, and *Methanosarcina acetivorans*) were also observed in which some enzymes a more limited capability for multiple nucleotide incorporation like the above mentioned TLPs even with tRNA substrates (Abad et al. 2010). Taken together, the diverse patterns of substrate specificity exhibited by individual TLP enzymes from different species is likely to be related to the unique functions for each of the enzymes, which have not yet been determined.

The consistent design of the RNA duplexes tested here (all containing the same 8-nt 3’-template sequence) enabled a more straightforward comparison of the enzymatic preferences of different enzymes from all domains of life than has been conducted previously. Using this assay, distinct biochemical behaviors of each enzyme in terms of binding and catalysis were readily observed (**Figures 2, 3, 4**). However, the simplicity of the standardized duplexes tested here leaves many questions about the boundaries of the 3’-5’ polymerase activity unanswered. Future studies with duplexes containing different template sequences and lengths will be important for understanding the limits of any potential applications of these enzymes. Although there were generally many similarities in the activity and behavior of AcaTLP2 and DdiTLP4 with the RNA duplexes tested here, there were also subtle differences, such as the propensity of AcaTLP2 to catalyze the addition of several nucleotides beyond what appears to be the full 5’-end of the extended duplex (Figure 2). In contrast, DdiTLP4 appears to be more likely to create a homogenous extended product that appears to correspond to addition of all 8 base pairs to the RNA substrate (Figure 2). The molecular basis for these differences is also unknown and may reflect different patterns of substrate recognition or distinct preferences related to the true biological substrates for these enzymes that have yet to be identified.

